# Vascular and neural transcriptomics reveal stage-dependent pathways to inflammation and cognitive dysfunction in a rat model of hypertension

**DOI:** 10.1101/2023.01.20.524921

**Authors:** Philipp Ulbrich, Lorena Morton, Michael Briese, Naomi Lämmlin, Hendrik Mattern, Md. Hasanuzzaman, Melina Westhues, Mahsima Khoshneviszadeh, Silke Appenzeller, Daniel Gündel, Magali Toussaint, Peter Brust, Torsten Kniess, Anja Oelschlegel, Jürgen Goldschmidt, Sven Meuth, Hans-Jochen Heinze, Grazyna Debska-Vielhaber, Stefan Vielhaber, Axel Becker, Alexander Dityatev, Solveig Jandke, Michael Sendtner, Ildiko Dunay, Stefanie Schreiber

**Affiliations:** Department of Neurology, Otto-von-Guericke University, Magdeburg, Germany; German Center for Neurodegenerative Diseases (DZNE) within the Helmholtz Association, Magdeburg, Germany; Institute of Inflammation and Neurodegeneration, Otto-von-Guericke University, Magdeburg, Germany; Institute of Clinical Neurobiology, University Hospital Würzburg, Würzburg, Germany; Department of Neurology, University Hospital Leipzig, Leipzig, Germany; Biomedical Magnetic Resonance, Faculty of Natural Sciences, Otto-von-Guericke University, Magdeburg, Germany; Comprehensive Cancer Center Mainfranken, University Hospital Würzburg, Würzburg, Germany; Helmholtz-Zentrum Dresden-Rossendorf, Institute of Radiopharmaceutical Cancer Research, Department of Neuroradiopharmaceuticals, Leipzig, Germany; Helmholtz-Zentrum Dresden-Rossendorf, Institute of Radiopharmaceutical Cancer Research, Department of Radiopharmaceutical and Chemical Biology, Dresden, Germany; Leibniz-Institute for Neurobiology, Magdeburg, Germany; Center for Behavioral Brain Sciences (CBBS), Magdeburg, Germany; Department of Neurology, Heinrich-Heine-University, Düsseldorf, Germany; Institute of Pharmacology and Toxicology, Otto-von-Guericke University, Magdeburg, Germany; Medical Faculty, Otto-von-Guericke University, Magdeburg, Germany

**Keywords:** hypertension, cerebral small vessel disease, SHRSP, RNA sequencing, FDG-PET

## Abstract

Chronic arterial hypertension causes cerebral microvascular dysfunction and doubles dementia risk in aging. However, cognitive health preservation by therapeutic blood pressure lowering alone is limited and depends on disease duration, the degree of irreversible tissue damage and whether microvascular function can be restored. This study aimed to understand molecular and cellular temporo-spatial pathomechanisms in the course of hypertension. We investigated the effects of initial, early chronic and late chronic hypertension in the frontal brain of rats by applying behavioral tests, histopathology, immunofluorescence, FACS, microvascular/neural tissue RNA sequencing as well as ^18^F-FDG PET imaging. Chronic hypertension caused frontal brain-specific behavioral deficits. Our results highlight stage-dependent responses to continuous microvascular stress and wounding by hypertension. Early responses included a fast recruitment of activated microglia to the blood vessels, immigration of peripheral immune cells, blood-brain-barrier leakage and an energy-demanding hypermetabolic state. Vascular adaptation mechanisms were observed in later stages and included angiogenesis and vessel wall strengthening by upregulation of cellular adhesion molecules and extracellular matrix. Additionally, we identified late chronic accumulation of Igfbp-5 in the brains of hypertensive rats, which is also a signature of Alzheimer’s dementia and attenuates protective Igf-1 signaling. Our study advances the knowledge of involved pathomechanisms and highlights the stage-dependent nature of hypertensive pathobiology. This groundwork might be helpful for basic and clinical research to identify stage-dependent markers in the human disease course, investigate stage-dependent interventions besides blood pressure lowering and better understand the relationship between poor vascular health and neurodegenerative diseases.

## Introduction

Midlife arterial hypertension is associated with cerebral small vessel disease (CSVD), accelerated cognitive decline and increased risk of all-cause dementia in aged individuals (Hazard Ratio (HR) 1.6), which even doubles, if hypertension persists into later life (HR 2.0) [1–5]. From a pathomechanistic point of view, chronic hypertension leads to cerebral microvascular dysfunction due to neurovascular uncoupling, decreased vascular reactivity and reduced cerebral blood flow, which impairs tissue supply with nutrients and oxygen and possibly brain waste clearance [2, 6–11]. Thereby hypertension represents a threat to healthy brain aging, but also leaves a time span of decades for prevention and therapy when recognized early. Despite increasing efforts in hypertension treatment and growing awareness of the deleterious effects of hypertension on cognitive function, only half of adult German or US hypertensive patients achieve adequate blood pressure (BP) control [6, 12], although it has been shown that BP reduction is able to lower mild cognitive impairment and dementia incidence [2, 13]. This points to a long-lasting modifiable therapeutic window for the prevention of cognitive decline in aging. At late stages, cognitive health preservation by BP reduction alone is however limited due to irreversible tissue damage and impaired microvascular function [5, 13]. Nonetheless, hypertension is still perceived as a disease state rather than a continuum with stage-dependent pathomechanisms. Current literature, which comprises experimental but also imaging-based human studies, suggests that hypertension-related pathobiology includes endothelial dysfunction, blood-brain-barrier (BBB) integrity loss, neuroinflammation, vascular wall remodeling and exacerbation of amyloid pathologies [3, 6, 14, 15]. Yet, a precise definition of molecular and cellular decision points in different stages of the disease course has not been sufficiently investigated, but is of pivotal importance for the development of targeted prevention and therapy. If fed a normal diet, non-transgenic spontaneously hypertensive stroke-prone rats (SHRSP), perfectly mimic this vulnerable hypertensive patient population in its initial and preclinical disease stages by combining arterial hypertension development and a polygenetic susceptibility to CSVD [16]. The frontal predilection of hypertensive pathology has been underlined by cognitive testing, magnetic resonance imaging (MRI) and a human microarray study in which the frontal cortex exhibited the most gene expression alterations [17–21]. Here, we investigated the frontal cortex of initial hypertensive (6-8 weeks) animals, that mimic newly diagnosed hypertension in midlife humans and of two older stroke-free age groups, that represent stages of early (24-25 weeks) and late chronic (32-34 weeks) hypertension. We provide a detailed characterization of molecular and cellular alterations at cerebral blood vessels and the surrounding neural tissue in response to different stages of arterial hypertension. Based on this groundwork future in-depth studies can select and manipulate cell type-specific perturbations to identify stage-dependent interventions next to blood pressure lowering for the preservation of healthy cognitive aging.

## Materials and Methods

### Animals

All experiments were approved by the Animal Care Committee of Saxony-Anhalt (reference number of license for animal testing 42502-2-1277 Uni MD). Male SHRSP (n = 75, Charles River Laboratories International Inc, Wilmington, Massachusetts, USA) and male Wistar control rats (n = 69, Charles River Laboratories, Research Models and Services, Germany GmbH, Sulzfeld, GER) aged 6-8 weeks, 24-25 weeks and 32-34 weeks were included in the study. All animals were housed with a natural light-dark cycle and had access to water and food *ad libitum*. The operators responsible for the experimental procedure and data analysis were blinded and unaware of group allocation throughout the experiments. Details on sample size in each experiment are shown in ***Supplementary Table 1***.

### Blood pressure measurement

Systolic BP was measured biweekly in Wistar rats and SHRSP (n = 3 per group) starting at week 8 until week 34 by the indirect tail-cuff method (BP-2000 Blood Pressure Analysis System, 4-channels, Visitech Systems, Apex, NC, USA). Pressure and pulse rate signals were continuously recorded and digitalized using BP-2000 Analysis Software. Systolic BP was determined as the mean of ten cuff inflation measurements.

### Open field test

CSVD patients exhibit altered gait, motor impairment and more subtle neuropsychiatric symptoms such as increased apathy [22, 23]. The open-field test was performed to determine the effect of hypertension on locomotor activity and anxiety-like behaviour in rats. On five consecutive days, the assessment took place in a square plexiglass box (1 m x 1 m) under low light conditions (30 lux). The rats were placed inside the box and allowed to move freely for 10 minutes while being recorded by an overhead camera. An automated tracking system (ActiMot, TSE Systems, Bad Homburg, GER) then analyzed activity, covered distance and time spent in the middle of the box. To standardise the olfactory background, 0.1% disinfectant Teta ExtraR (Fresenius Germany, containing 2.5% polyhexanide and 8% didecyldimethylammonium) was used for cleaning before the first and after each test.

### Prepulse inhibition test

Prepulse inhibition (PPI) reflects sensorimotor gating and is involved in attentional processing, mainly controlled by the frontal cortex, including the anterior cingulate, medial prefrontal and motor areas [24–26]. Frontal brain regions are frequently affected by CSVD pathology and were linked to a decline in focused attention and processing speed [27]. The PPI test is carried out to examine, whether the acoustic startle response (ASR), a sudden whole-body movement, is mitigated by an acoustic prepulse. The animals were tested in two ventilated sound-attenuating startle chambers (startle response box, TSE Systems, Bad Homburg, Germany). One rat from the control group and one SHRSP were tested simultaneously. Acoustic stimuli were delivered via one speaker on each site of the startle chamber mounted at a distance of 4 cm. Strength of movement (expressed in grams) to an acoustic stimulus was measured by a force transducer (0–10 N) and was defined as the average of fifty 2-ms readings collected from stimulus onset. A background noise level of 68 dB was maintained throughout the test session. After a habituation period of 5 min, five startle pulses (30 ms, 120 dB) were presented to measure the ASR. Afterwards, PPI was measured. The prepulses were acoustic stimuli of 72, 76, 80, or 84 dB of 20 ms in duration. The interval between the prepulse and pulse was 80 ms. The session consisted of six presentations of startle pulses, prepulses alone at four intensities, prepulses at any intensity followed by startle pulses, no stimulus trials, in randomised order, and finally five startle pulses. The interval between two trials ranged from 11 to 19 s. One session lasted for about 25 min. For no stimulus trials, readings were recorded over periods when no stimulus was presented to assess gross motor activity during the test session. PPI was defined as the percentage reduction of startle magnitude in the presence of the prepulse compared to the magnitude in the absence of the prepulse (100 - (100 - magnitude on prepulse + pulse trial/magnitude on pulse trial)).

### Brain tissue collection and preparation for histopathology and immunofluorescence

For anaesthesia, pentobarbital (40 mg/kg body weight) was injected intraperitoneally in all animals. Transcardial perfusion was conducted with 120 mL of phosphate buffered saline (PBS) followed by fixation with 120 mL of 4% paraformaldehyde (PFA) within 8 min. Brains were removed, fixed in 4% PFA for 48 h, cryoprotected in 30% sucrose for 6 days and frozen in methylbutane at −80 °C. Using a cryostat, the brains were then sliced from the frontal to the occipital pole in 30 µm thick sections.

### Histopathological analysis

Histological hematoxylin and eosin (HE) staining enables to study hallmarks of microvascular pathology, including cerebral microbleeds (CMB) and enlarged perivascular spaces (EPVS). For histological analysis of EPVS n = 5 frontal coronal slices with both hemispheres were taken per animal in all groups. Brain slices were washed twice with distilled water, incubated with haematoxylin (Carl Roth GmbH + Co. KG, Karlsruhe, GER), washed again, followed by bluing under running tap water and another distilled water rinse. Next, staining was performed with 1% eosin solution (Carl Roth GmbH + Co. KG, Karlsruhe, GER) for 40 s. After dehydration with increasing concentrations of Rotisol (Carl Roth GmbH + Co. KG, Karlsruhe, GER) the tissue was embedded in Xylene (Carl Roth GmbH + Co. KG, Karlsruhe, GER) and mounted with coverslips using Histomount (Fisher Scientific GmbH, Schwerte, GER). HE-stained slices were subjected to light microscopic examinations (Leica DMR). To obtain the frequency of EPVS in the frontal cortex, we calculated the number of small vessel segments with EPVS / total number of small vessel segments per field of view (FOV). Small vessel segments included parenchymal arterioles and venules with a diameter greater than 15 µm. EPVS were identified as unstained, distended perivascular areas. We analysed 30 cortical FOVs per animal in all groups with a 20x objective. Additionally, we randomly selected 10 EPVS per animal that ran along the plane of the slice and quantified the EPVS area in 34-week-old Wistar rats and SHRSP using a polygon tool in ImageJ software and a 40x objective.

### Immunofluorescence staining

Immunofluorescence staining of multiple target proteins was used to investigate the spatial interaction of glial cells, blood vessels and areas of BBB leakage. The staining was performed as previously described [28]. In short, for each staining, two coronal brain slices with both hemispheres were stained per animal. They were repeatedly washed in PBS, blocked with 10% donkey serum/0.5% TritonX (Sigma, St Louis, MO, USA) and incubated with primary antibodies overnight at 4°C (IBA1 - ionized calcium-binding adapter molecule 1, microglial marker; GFAP - glial fibrillary acidic protein, astrocytic marker; IgG - immunoglobulin G, BBB breakdown marker; STL - solanum tuberosum lectin, endothelial marker). The following day they were repeatedly washed again, incubated with secondary reagents for two hours and mounted on slides with Fluoromount Aqueos Mounting Medium (Merck, F4680). Used primary and secondary reagents and dilutions are listed in ***Supplementary Table 2***.

### Image acquisition and analysis

Immunofluorescence images were acquired using a Zeiss confocal microscope (LSM 700) and EC Plan-Neofluar 20×/0.50 M27 objective. For each antibody, the gain and laser power were kept equal for all animals. Three frontal cortical regions, which are involved in PPI and where BBB leakage was most prominent were analysed. Therefore, in each slice we acquired two overview z-stack images (8-bit, 512 × 512 pixels with pixel size of 1.25 × 1.25 µm^2^) in the motor, anterior cingulate and medial prefrontal cortex, respectively, resulting in 12 cortical FOVs per animal. Image analysis was performed using openly available ImageJ software and Python3-based in-house routine. Pre-processing included computation of z-projection (maximum intensity) and subtraction of background (rolling ball radius: 50 pixels) and was performed in ImageJ. For microglia soma quantification and vessel segmentation, additionally, the median filter was applied to remove “salt and pepper” noise. Microglia activation has been linked to acute BBB breakdown and it is widely accepted that there are clear morphological changes in microglia after activation, such as enlargement of soma and a reduction of microglial branches [29–31]. For the evaluation of microglia morphology, two thresholding and size exclusion criteria were applied to measure the mean cellular branching (thresholding 1: autothresholding with triangle dark option, > 250 µm^2^) and soma area (thresholding 2: autothresholding with moments dark option, > 50 µm^2^; example in ***Supplementary Figure 2***). GFAP is a well-established astrocytic activation marker and IgG is only able to diffuse into the vessel wall and brain parenchyma, when BBB integrity is compromised [32]. Since GFAP expression and IgG levels highly varied between the age groups, fixed threshold values were applied to assess astrocyte activation and BBB breakdown (for both: threshold 25). To avoid wrongly segmented pixels or incomplete cells, size exclusion criteria was set for IgG > 5 µm^2^ and GFAP > 30 µm^2^ [29]. The positive area was quantified and expressed as % IgG area per FOV, mean microglial soma/branching area and mean GFAP cluster area. Microglia soma and GFAP+ clusters were counted as previously described and normalized to an area of 1 mm^2^ [29]. For vessel segmentation in STL-labelled images, the OMELETTE framework was adapted [33] (https://gitlab.com/hmattern/omelette). First, blood vessels were enhanced using a multi-scale Frangi filter [34]. The filter’s vessel sensitivity parameter gamma was selected empirically to be 40% of the maximum absolute Hessian eigenvalues so that a multi-scale Frangi filter with an image-specific gamma was applied (filter scales 1, 2, 3, 4, 5 pixels). Second, the blood vessels were segmented from the enhanced images by applying a hysteresis thresholding [35]. Three-class Otsu’s method was used to estimate the two required thresholds from the enhancement distribution (self-tuned threshold estimation). Last, a centreline of the segmented vasculature was generated using skeletonization [36]. The total length of the segmented microvasculature was approximated as the sum of all centreline pixels multiplied by the pixel edge length and normalized to an area of 1 mm^2^.

To assess whether glial cells are recruited to the blood vessels, the relative number of microglia soma and GFAP+ clusters tangent to the vessel segmentation was estimated. In late chronic hypertension, additionally, blood vessel-associated microglia were manually differentiated as located at vascular junctions and at non-junctional vessel segments, since it was recently shown, that they can control blood flow of the downstream vascular bed [37]. To elucidate the spatial relation of glial cells to BBB breakdown and thereby the reactivity and migration pattern in response to vascular injuries, the recently introduced vessel distance mapping [38–40] was adapted. For each segmentation individually (i.e., microglia soma, GFAP clusters, BBB breakdown), the Euclidian distance to the closest segmented cluster of interest was computed [41]. To measure the shortest distance from IgG+ pixels to GFAP+ clusters or microglia soma, the values from IgG distance maps were extracted at the location of GFAP+ clusters and microglia soma. Group distributions were compared by cumulative distribution function (CDF) plots.

### FACS analysis of cortical leukocytes and endothelial cells

Fluorescence-activated cell sorting (FACS) analysis was used to quantify numbers of infiltrating peripheral immune cells and of endothelial cells in the entire cortex. After transcardial perfusion with PBS containing EDTA, brains were dissected and divided into two sagittal halves, of which one was processed for FACS analysis and the other one for microvessels isolation. For flow cytometric analysis, a single-cell suspension was created. The cerebral cortex was cleaned of meninges and the entire cell populations were isolated by enzymatic digestion and mechanical dissociation using the Multi Tissue Dissociation Kit 1 (Miltenyi Biotec, Bergisch Gladbach, GER) according to manufacturer instructions followed by a 30% density gradient centrifugation. After myelin aspiration, cells were filtered using a 70 µm cell strainer, washed with RPMI and resuspended in FACS buffer (2% fetal calf serum in PBS). Cells were incubated for 15 min at 4 ºC with a purified mouse anti-rat CD32 FcγII antibody (clone D34-485) to block unspecific binding. Thereafter, cells were stained with a mixture of fluorochrome-conjugated monoclonal antibodies in 100 µL of FACS buffer for 30 min at 4ºC. Cells were washed twice and resuspended in 200 µL of FACS buffer. A mixture of antibodies included Zombie NIR™, anti-rat CD45 (OX-1), anti-rat CD31 (PECAM-1) and anti-rat Ki-67 (REA1123). Optimization was performed using antibody titrations and Fluorescence Minus One controls to assess background fluorescence in the respective detection channel. Samples were acquired on Attune NxT Flow Cytometer (ThermoFisher Scientific, Waltham, MA, USA) and analyzed with Flowjo Analysis Software v10.5.3.

### Cortical microvessel isolation and RT-qPCR

Microvessels were additionally isolated from the entire cerebral cortex to validate and generalize findings from experiments in the frontal cortex. The remaining second cortical hemisphere, which was not utilized for FACS analysis, was used to obtain pure cortical microvessels (modified and optimized from a recently described protocol [42]). Briefly, cerebral cortices from late chronic hypertensive rats and age-matched controls were cleaned from the meninges, cerebellum, deep brain structures and residual white matter. The tissue was homogenized using a 2 mL Dounce tissue grinder. The homogenate was transferred into a 15 mL tube, centrifuged and resuspended in 15% (wt/vol) dextran solution-DPBS. Samples were further centrifuged at 10,000g for 15 min. The resulting myelin layer together with the supernatant was carefully removed and the microvessel-containing pellet was transferred through a 40 µm cell strainer for microvessel collection. Thereafter, the cell strainer was reversed and washed into a new 50 mL tube for the retrieval of microvessels. The cell suspension was centrifuged at 400g for 10 min at 4 ºC and the purity of the microvessels was quickly evaluated mounting 20 µL of microvessels suspension and observed under an inverted microscope. Brain debris, myelin and white matter were not observed in the final preparation. Collected microvessels were pelleted at 20,000g for 10 min and resuspended in 350 µL of RLT Plus Buffer from the RNeasy Micro Kit. Total RNA was isolated using the RNeasy Micro Kit according to the manufacturer’s instructions. Samples were dissolved in an appropriate amount of RNAase-free ddH_2_O and RNA concentration and purity were determined using NanoDrop 2000 (ThermoFisher Scientific, Waltham, MA, USA). Levels of *Vegfb* and *Igfbp-5* gene expression were quantified by reverse transcription and quantitative RT-PCR. Relative gene expression was determined using the TaqMan® RNA-to-CT™ 1-Step Kit (ThermoFisher Scientific, Waltham, MA, USA). Reactions were developed in a LightCycler® 96 (Roche, Basel, Switzerland). Reverse transcription was performed for 30 min at 48 ºC followed by 10 min at 95 ºC. Subsequently, a two-step amplification was run for 55 cycles, comprising denaturation for 15 s at 95 ºC and annealing/elongation for 1 min at 60 ºC. TaqMan® Gene Expression Assays (ThermoFisher Scientific, Waltham, MA, USA) were used for mRNA amplification of *GAPDH* (Rn01775763_g1), *Vegfb* (Rn01454585_g1) and *Igfbp-5* (Rn00563116_m1). *GAPDH* mRNA expression was chosen as a reference for normalization and target/reference ratios were calculated with the LightCycler ® 96 Software release 1.1 (Roche, Basel, Switzerland).

### Laser microdissection and RNA isolation for RNA sequencing

To evaluate stage-dependent gene expression alterations in cerebral blood vessels and neural tissue separately, the different structural compartments of the brain parenchyma were isolated via laser microdissection and utilized for RNA sequencing (RNAseq). Brains were embedded in TissueTek and frozen with liquid nitrogen. Until further processing, the brain tissue was stored at -80°C to preserve the RNA. Histological sections were prepared with a cryotome (Leica CM1950) at -20°C. As soon as the frontal pole became visible, the first 990 µm of the brain tissue was discarded. Subsequently, n = 16 brain slices per animal were put on PEN membrane slides (four 15 µm-thick slices per slide, 90 µm discarded after every four sections) and immediately prepared for laser microdissection. In detail, slides were washed in 70% ethanol for 20 s and dried at 40 °C for 10 min to preserve high RNA quality. Slides were then placed in toluidine blue for 3 min, washed four times in DEPC water for 20 s, incubated in 70% ethanol for 3 min and dried for 10 min at 40 °C. Each slide was individually placed in a Falcon tube filled with silica gel and stored on ice. During laser microdissection (Leica LMD 6000) in the frontal cortex, approximately 250 randomly chosen small vessels with a luminal diameter ranging from 10-80 µm (corresponding to parenchymal capillaries, arterioles and venules [43]) and four neuron clusters (each per brain section) were dissected per slice. Thus, approximately 4000 small vessels and 64 neuronal clusters were collected per animal and separated as blood vessel-enriched and neuron-enriched (neural) tissue samples. RNA was then extracted using the PicoPure RNA Isolation Kit (ThermoFisher Scientific, Waltham, MA, USA). After washing away all impurities, the RNA was eluted and the pure RNA was stored at -80 °C.

### RNA sequencing analysis

RNA-seq libraries were prepared using the QuantSeq 3’ mRNA-Seq Library Prep Kit FWD (Lexogen). Data analysis was performed as recommended by Lexogen’s QuantSeq Data Analysis Pipeline User Guide. Briefly, sequencing read quality was assessed with FastQC v0.11.5 (https://www.bioinformatics.babraham.ac.uk/projects/fastqc/) and reads were trimmed with the bbduk from BBTools v37.93 (https://jgi.doe.gov/data-and-tools/bbtools/). Reads were aligned to the ENSEMBL Rattus norwegicus (Rnor6) reference genome with STAR v2.5.3a [44] and counts per gene were quantified with HTSeq-count v0.6.1 [45] using the ENSEMBL Rattus norwegicus (Rnor6) annotation. Differential expression analysis was performed with DESeq2 v1.18.1 [46]. The sequencing files are available at NCBI’s Gene Expression Omnibus [47] accession number GSEXXXXXX.

### RT-qPCR validation

To validate differentially expressed genes, which were identified via RNAseq we performed RT-qPCR for selected target genes. RNA generated for RNAseq was reverse-transcribed with random hexamers using the First Strand cDNA Synthesis Kit (ThermoFisher Scientific, Waltham, MA, USA). Reverse transcription reactions were diluted 1:5 in water. Reactions were set up with Luminaris HiGreen qPCR Master Mix (ThermoFisher Scientific, Waltham, MA, USA) on a LightCycler® 96 thermal cycler (Roche, Basel, Switzerland). Primers are listed in ***Supplementary Table 3***. The following RNAseq results were confirmed by RT-qPCR: Blood vessels - *RT1* (up at 25 and 34 weeks), *Serping1* (up at 25 weeks) *Igfbp-5, Nov, Ctgf, Tgfa, Cdh5* (up at 34 weeks). In neural tissue *Pld1, Nppa* (down at 8, 25 and 34 weeks), *Atp11b, Zfp462* (down at 34 weeks) and *RT1* (up at 24 weeks) were similarly dysregulated (***Supplementary Figure 1***).

### ^18^F-FDG-PET

Gene expression alterations in the hypertensive rat brain suggested an upregulation of energy-demanding processes. Using ^18^F-fluorodeoxyglucose positron emission tomography (^18^F-FDG-PET) imaging, which measures brain glucose consumption, we assessed whether these molecular and cellular changes affect brain metabolism and are transferable to further brain regions. In all three age groups SHRSP and Wistar rats were injected i.p. with ^18^F-FDG (1 to 2 mL, diluted in saline). The youngest age group received 21.7 ± 2.5 MBq ^18^F-FDG, for the other two age groups the dose was increased to 43.2 ± 4.2 MBq for better signal-to-noise ratios. Animals were injected in the awake state and remained in their home cages after injection. After 30 min animals were anaesthetized with 2% isoflurane in 60% oxygen and placed on a thermostatically heated animal bed. PET-scanning in a small-animal PET/MR (Nanoscan, Mediso, Budapest, HUN) started 45 min after ^18^F-FDG injection. Static PET scans of the head were performed followed by T1-weighted GRE MR imaging for anatomical correlations and attenuation correction. PET acquisition time was 15 min. List mode data from PET scans were reconstructed by Ordered Subset Expectation Maximization (OSEM3D) with four iterations, six subsets and a voxel size of 0.4 mm isotropic (Nucline v2.01, Mediso, HUN). Hematocrit (0.5 to 0.64) and blood glucose concentration (5.7 to 8.6 mmol/L, Accu-Chek Performa Nano, Roche, Mannheim, GER) were measured shortly before injecting the radiotracer.

### Data analysis and statistics

Mean values per animal were entered into statistical analysis. Gaussian distribution of the data points was verified using the Shapiro-Wilk test. For multiple comparisons, two-way analysis of variance (ANOVA) followed by Holm-Šidák’s post hoc test with group (SHRSP vs. Wistar) and age as categorical variables was applied to test statistical significance between hypertensive and control rats and to reveal age-related effects. This was done for BP measurements, behavioral tests, EPVS frequency, immunofluorescence experiments and leukocyte FACS. For two group comparisons, the statistical significance was analyzed using a two-tailed Student’s t-test. This included the EPVS area, endothelial FACS, Vegfb and Igfbp-5 expression in isolated microvessels. One-way ANOVA with Holm-Šidák’s correction for multiple comparisons was applied for Igfbp-5 expression in laser-dissected blood vessels. CMB prevalence probability was analyzed in cumulative data from previous publications [28, 48, 49], containing in total n = 96 Wistar rats and n = 203 SHRSP aged 8 to 44 weeks. Age-dependent prevalence probability was visualized in each strain with simple logistic regression and the age and hypertension effects on CMB prevalence were estimated using the multivariate logistic regression (details in ***Supplementary Table 4***). Group differences in CDF plots were assessed statistically using a two-sample, two-sided Kolmogorov-Smirnov (K-S) test, which was used for the shortest IgG – glial cell distances. All analyses were performed using GraphPad Prism software and Python3. STRING interaction networks were created for each dataset from all differently expressed genes. Interaction sources were experiments, databases, co-expression, neighborhood, gene fusion and co-occurrences. Gene interactions were only shown with high (interaction score > 0.7, for blood vessels) and highest confidence (interaction score > 0.9, for neural tissue) (www.string-db.org). For gene ontology (GO) and KEGG (Kyoto Encyclopedia of Genes and Genomes) enrichment analysis we used the Database for Annotation, Visualization and Integrated Discovery (DAVID) (https://david.ncifcrf.gov/). Background datasets were selected separately for blood vessels (early chronic hypertension: 11.928 transcripts; late chronic hypertension: 13.984 transcripts) and neural tissue (early chronic hypertension: 13.610 transcripts; late chronic hypertension: 13.821 transcripts), including genes with mean transcript number > 1. Significance for GO-term enrichments was set using the EASE score with a p-value < 0.05. To compare single transcripts between the four datasets, counts were normalized to the respective total transcript number of the dataset. For individual cell-specific genes, we analyzed tissue enrichment by applying a paired, two-tailed Student’s t-test (see Results – “Validation of method”). ^18^F-FDG PET-data were spatially aligned to a Wistar MR template [50] using the MPI-Tool™ software (Advanced Tomo Vision, GER). After alignment, data-files were converted to 32-bit TIFF-files for processing in FIJI and MATLAB™ (v. R2017b). In FIJI, brain-data were cut out of the PET-data and intensity-normalized. Voxel-wise unpaired two-tailed heteroscedastic t-tests were performed in MATLAB™. Files of group-mean ^18^F-distributions were calculated in FIJI by adding in each group individual brain data and dividing by the number of animals. Data were converted to DICOM-files and rendered in the DICOM-viewer Osirix MD™ (v. 13.0.0, Pixmeo, CH).

## Results

### Blood pressure measurement

SHRSP exhibited elevated systolic BP from the age of 8 weeks onwards compared to age-matched Wistar rats (group main effect: p < 0.0001). Systolic BP significantly increased with age and plateaued in chronic hypertension (8 vs. 24 weeks p = 0.01; ***Figure 1a***).

**Figure 1.**
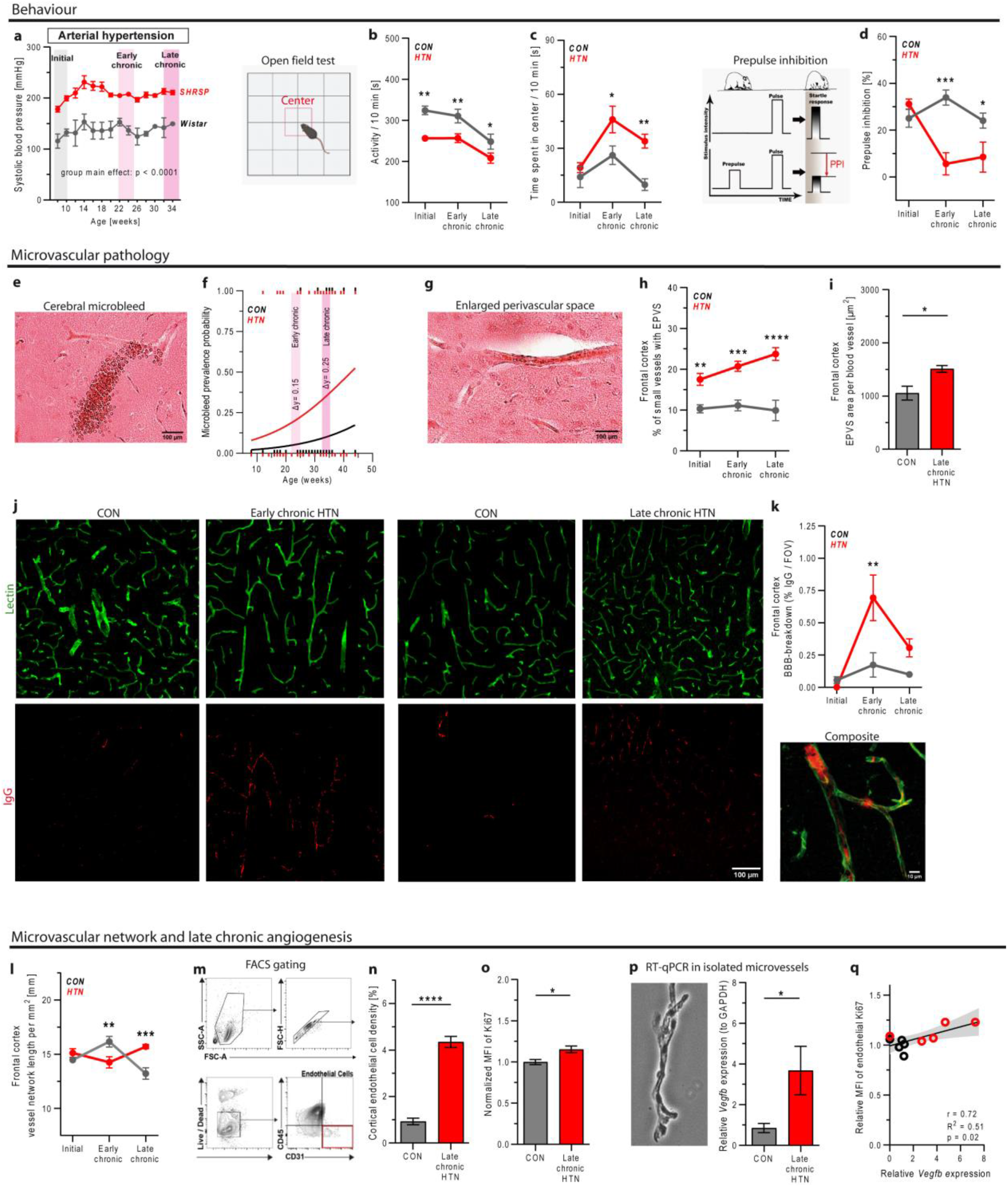
Hypertension causes behavioral alterations and microvascular pathology in the frontal cortex of SHRSP. (**a-d**) Behavior experiments: (**a**) Longitudinal, biweekly systolic blood pressure measurements in Wistar rats and SHRSP (n = 3 per group). In initial, early chronic and late chronic hypertension and respective controls (n = 10 per subgroup) we performed an open field test (**b**,**c**) to analyze overall activity and time spent in the center of the arena followed by a prepulse inhibition test (**d**). (**e**,**f**) Cerebral microbleed prevalence was examined in the entire brain of HE-stained slices and is shown as a logistic regression model with age as the independent variable. This includes n = 203 hypertensive and n = 96 normotensive rats, aged 8 to 44 weeks from cumulative data of previous publications [27, 51, 52]. Group differences are highlighted for early and late chronic hypertension and expressed as Δy. (**g-i**) Relative number of small blood vessels with EPVS and EPVS area per blood vessel were analyzed in the frontal cortex of HE-stained brain slices. (**j**,**k**) BBB breakdown was assessed using immunostaining of endothelial cells and IgG plasma protein and was quantified as %IgG per FOV. (**j, l**) Cortical microvascular network length in the frontal cortex. (**m-o**) FACS analysis of cortical endothelial cells in late chronic hypertension and age-matched controls: (**m**) gating strategy, (**n**) endothelial cell density and (**o**) their mean fluorescence intensity of Ki67, a mitotic cell marker. (**p**) Using RT-qPCR Vegfb expression level was analyzed in isolated cortical microvessels (an exemplary image is shown by inverted microscopy). (**q**) Pearson’s correlation between angiogenic Vegfb expression and endothelial Ki67 proliferation marker MFI. In **h, i, k, l** and **n-p** each of subgroups comprised n = 5 or 6 animals. Two-way ANOVA and Holm-Sidák’s post hoc test for multiple comparisons were applied to test for significant differences in **a-d, h, k** and **l**. Two-tailed Student’s t-test was applied in **i** and **n-p**. Significant group differences are highlighted via * p < 0.05; ** p < 0.01; *** p < 0.001, **** p < 0.0001. Data are presented as mean ± s.e.m. Abbreviations: BBB, blood-brain barrier; CD31, cluster of differentiation 31, endothelial marker; CD45, cluster of differentiation 45, leukocyte antigen; CON, control; EPVS, enlarged perivascular spaces; FACS, fluorescence-activated cell sorting; FOV, field of view; FSC-A and FSC-H, forward scatter and cell size marker; HE, hematoxylin and eosin; HTN, hypertension; IgG, Immunoglobulin G; min, minute; Ki67, marker of proliferation Ki-67; MFI, mean fluorescence intensity; PPI, prepulse inhibition; RT-qPCR, reverse transcription quantitative real-time PCR; SHRSP, spontaneously hypertensive stroke-prone rat; SSC-A, side scatter, cell granularity marker.

### Open field test

In order to identify onset and severity of cognitive alterations in our cohort of hypertensive rats, we compared animals cross-sectionally in initial, early chronic and late chronic hypertension to age-matched normotensive controls. Locomotor activity was already significantly attenuated in initial hypertension (adj. p = 0.002), which persisted in early chronic (adj. p = 0.009) and late chronic hypertension (adj. p = 0.04; ***Figure 1b***), possibly indicating motivational disorder or apathy-like behavior. Time spent in the center square was not altered in initial hypertension but significantly increased in both chronic hypertensive stages (for both adj. p = 0.004; ***Figure 1c***), suggesting reduced anxiety.

### Prepulse inhibition test

Recent studies have shown that alterations in PPI are a sensitive measure for dysfunction of neuronal circuits in the forebrain [51, 52]. Such alterations have been observed models of frontotemporal dementia [53, 54] and AD [55]. In order to find out whether similar alterations are detectable hypertensive rats, we performed analyses of a PPI test. PPI was unchanged in initial hypertension but steeply sloped in early and late chronic hypertension compared to controls (early chronic: adj. p < 0.0001; late chronic: adj. p = 0.02; ***Figure 1d***). PPI is connected to attentional processing and a cognitive limitation in chronic hypertensive rats.

### Microvascular pathology in the frontal cortex

Hypertensive microvascular pathology comprises CMB formation, PVS enlargement and BBB integrity loss [56]. CMB prevalence probability was overall tripled in SHRSP compared to Wistar rats (OR 3.2, p = 0.0007). The probability difference raised from early (group difference Δy = 0.15) to late chronic hypertension (Δy = 0.20), showing that the longer hypertension persisted in a chronic stage the more likely microvascular pathologies occurred compared to normotensive controls. There were no CMB in initial hypertensive animals and age-matched controls < 10 weeks of age (***Figure 1e, f and Supplementary Table 4***). Contrary, EPVS in the frontal cortex were already more frequent in initial hypertension compared to age-matched controls (adj. p = 0.004), further intensifying in chronic hypertensive stages (early chronic: adj. p = 0.0005; late chronic: adj. p < 0.0001; ***Figure 1g, h***). In late chronic hypertension, the area of EPVS per blood vessel was significantly increased by around 50% compared to age-matched controls (p = 0.01; ***Figure 1i***). Plasma protein leakage as a result of BBB integrity loss was absent in normotensive and hypertensive 8 weeks old animals but occurred in chronic hypertension and the corresponding age-matched controls. There was a significant increase in early chronic hypertensive animals compared to controls (adj. p = 0.002), which declined again in late chronic hypertension (***Figure 1j, k***). Immunostaining of endothelial cells was used to characterize the microvascular network. In initial hypertension vascular network length was unaltered. With age vascular network length increased in late chronic hypertensive animals, resulting in group differences of around 20% (adj. p = 0.0008; ***Figure 1l***). FACS analysis of cortical endothelial cells and qPCR of isolated microvessels validated active angiogenesis in late chronic hypertension, showing increased endothelial cell density (p < 0.0001), elevated proliferation marker Ki67 (p = 0.02) and angiogenic *Vegfb* expression (p = 0.048, ***Figure 1m-p***). Microvascular *Vegfb* expression positively correlated with endothelial Ki67 levels (r = 0.72, p = 0.02, ***Figure 1q***), suggesting that the amounts of angiogenic factors may determine endothelial proliferation rates.

### Immune cell infiltration and glial cell reactivity in chronic hypertension

In early chronic hypertension, when BBB breakdown was most severe, FACS analysis revealed significantly increased cortical infiltration of peripheral leukocytes (adj. p = 0.04). With age, it increased in both groups, but was not elevated anymore in late chronic hypertension (***Figure 2a, b***). Immunostaining of IBA1+ microglia and GFAP+ astrocytes was used to assess glial activation in the frontal cortex, estimating cell density, mean IBA1+ soma/branching area and mean area of GFAP+ clusters. Already in response to initial hypertension and in the absence of BBB breakdown microglia showed an activated phenotype with significantly increased soma size (adj. p = 0.04), while branching area and astrocytic markers were unchanged. In early chronic hypertension, when BBB breakdown was most severe, microglia showed no morphological features of activation and GFAP+ astrocyte density was even decreased (adj. p = 0.0009) compared to age-matched controls. In late chronic hypertensive animals, activated microglia presented again with significantly increased soma (adj. p = 0.02) and branching area (adj. p = 0.001), despite the decline in BBB-breakdown. No differences were noticed in astrocyte GFAP expression (***Figure 2c-e and g-i, Supplementary Figure 2***). This suggests that glial activation in chronic hypertension is not a precise response to the severity of BBB breakdown. Correlation analysis revealed that in response to increasing BBB-breakdown severity, microglia retracted their branches (controls: r = -0.65, p = 0.04; chronic hypertension: r = -0.78, p = 0.004) and GFAP+ astrocyte density increased (controls: r = 0.66, p = 0.04; chronic hypertension: r = 0.64, p = 0.03) similarly in chronic hypertension and age-matched controls, but reactive soma enlargement (r = 0.87, p = 0.001) and the increase of GFAP+ cluster area (r = 0.76, p = 0.01), was only visible in controls and not in chronic hypertension (***Figure 2f, Supplementary Figure 2***). Therefore, glial reactivity via microglial morphology changes and astrocytic GFAP expression is a response to BBB breakdown in normotensive rats but is altered or reduced in chronic hypertension. To determine spatial migration patterns of glial cells in response to BBB-breakdown and whether they were drawn to the site of injury, we next measured the distance from each IgG positive pixel to the closest IBA1+ microglia and GFAP+ astrocyte. GFAP+ astrocytes were found more distant to BBB breakdown in early and late chronic hypertension compared to age-matched controls (for both p < 0.0001, K-S test, ***Figure 2j, k***). There were no distance differences between the groups for BBB-breakdown and microglia. Density of microglia was overall unchanged but declined with age in both groups (***Supplementary Figure 2***). Next, we analyzed the portion of glial cells associated with blood vessels, showing that microglia were more frequently associated with blood vessels already in initial (adj. p = 0.0001) and persistently in early (adj. p = 0.005) and late chronic hypertension (adj. p = 0.01, ***Figure 2l, m***). In late chronic hypertension, this increase of blood vessel-associated microglia was made up of cells at vascular junctions (p = 0.02), whereas the portion at non-junctional vessel segments was unchanged (***Figure 2n, o***). Thus, we conclude, that microglial recruitment to blood vessels is a fast and persistent response to hypertension and may be the reason why microglia are located in the vicinity of vascular injury. The portion of blood vessel-associated GFAP+ astrocytes was unchanged between groups (***Supplementary Figure 2***), suggesting that the number of reactive perivascular GFAP+ astrocytes is unchanged, but particularly at sites of vascular injury, reactive astrocytic GFAP expression is reduced in chronic hypertension.

**Figure 2.**
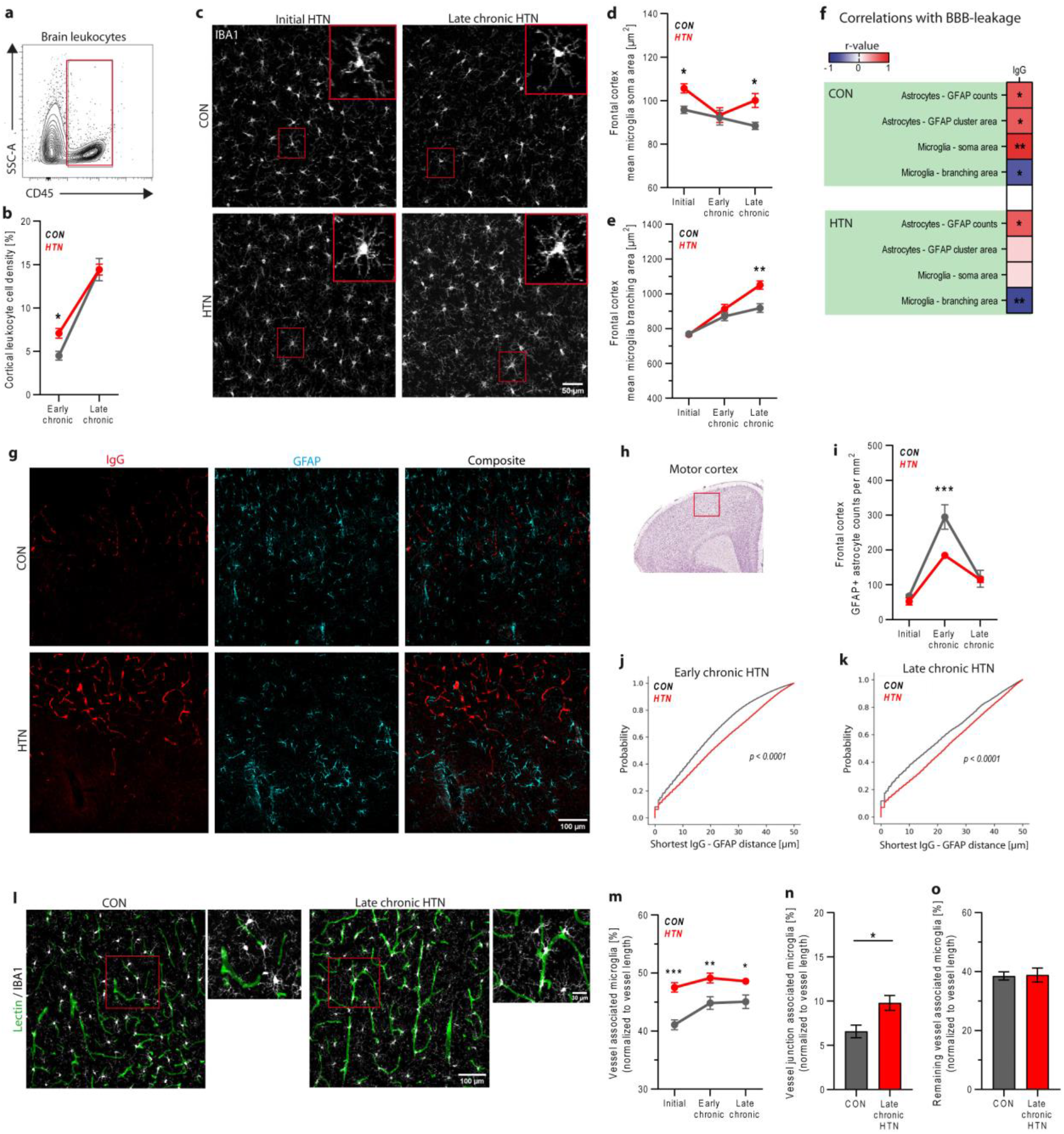
Hypertension causes peripheral leukocyte infiltration and glial activation in the frontal cortex of SHRSP. (**a**,**b**) Infiltrating blood leukocyte density was quantified via FACS in early and late chronic hypertension and age-matched controls. Markers of glial cell activation were analyzed in initial, early chronic and late chronic hypertension and respective controls using immunostaining of microglial IBA1 and astrocytic GFAP in the frontal cortex. (**c-e**) Microglia morphology was analyzed by measuring mean cellular soma and branching area. (**f**) Pearson’s correlation between glial activation markers and BBB breakdown was analyzed in both chronic hypertensive groups and respective controls, using mean values per animal. The correlation coefficient is color-coded with a double gradient, in which blue represents negative and red positive correlations, *p < 0.05, **p < 0.01. (**g**,**h**) Exemplary FOVs in the motor cortex show strong BBB-breakdown and reactive astrocytic GFAP expression. (**i**) Number of GFAP+ astrocytes per mm^2^ was quantified and (**j**,**k**) the shortest distance from BBB-breakdown (IgG positive pixel) to the closest GFAP+ astrocyte was estimated and displayed as cumulative relative frequency. (**l**) Exemplary FOVs in late chronic hypertension and an age-matched control show the interaction of blood vessels and microglia. (**m**) Percentage of microglia, associated with blood vessels was quantified in the frontal cortex. (**n**,**o**) Additionally, the percentages of microglia, associated with vascular junctions and non-junctional vessel segments were quantified in late chronic hypertensive animals and age-matched controls. In **b, d, e, i-k** and **m-o** each subgroup comprised n = 5 or 6 animals. Two-way ANOVA and Holm-Šidák’s post hoc test for multiple comparisons were applied to test for significant differences in **b, d, e, i** and **m**. Two-tailed Student’s t-test was applied in **n, o** and K-S test in **j, k**. Significant group differences are highlighted via * p < 0.05; ** p < 0.01; *** p < 0.001. Data are presented as mean ± s.e.m. Abbreviations: BBB, blood-brain barrier; CD45, cluster of differentiation 45, leukocyte antigen; CON, control; FACS, fluorescence-activated cell sorting; FOV, field of view; GFAP, glial fibrillary acidic protein, astrocyte marker; HTN, hypertension; IBA1, ionized calcium-binding adapter molecule 1, microglia marker; IgG, Immunoglobulin G, BBB leakage marker; SSC-A, side scatter, cell granularity marker.

### Validation of vascular and neural microdissection and RNA sequencing

To identify stage-dependent transcriptomic signatures in chronic hypertension, we dissected blood vessels and neural parenchyma (***Figure 3a***), performed RNAseq and analyzed differentially expressed genes. We identified significantly higher transcript numbers of vascular cell-specific genes within the dissected blood vessels compared to the neural tissue. These included genes characteristic for endothelial cells (*Cldn5, Abcba1, Vwf*; ***Figure 3b***), pericytes (*Pdgfrb, Cspg4*), vascular smooth muscle cells (*SMC*) (*Acta2, Tpm1, Tagln*), perivascular astrocytes (*Aqp4*), fibroblast-like brain cells (FB) (*Col1a1, Apod, Mgp*) and lymphatic marker-positive cells (*Lyve1, Flt4*) (***Supplementary Table 5***). The laser microdissection was instrumental to isolate all core structures of the neurovascular unit.

**Figure 3.**
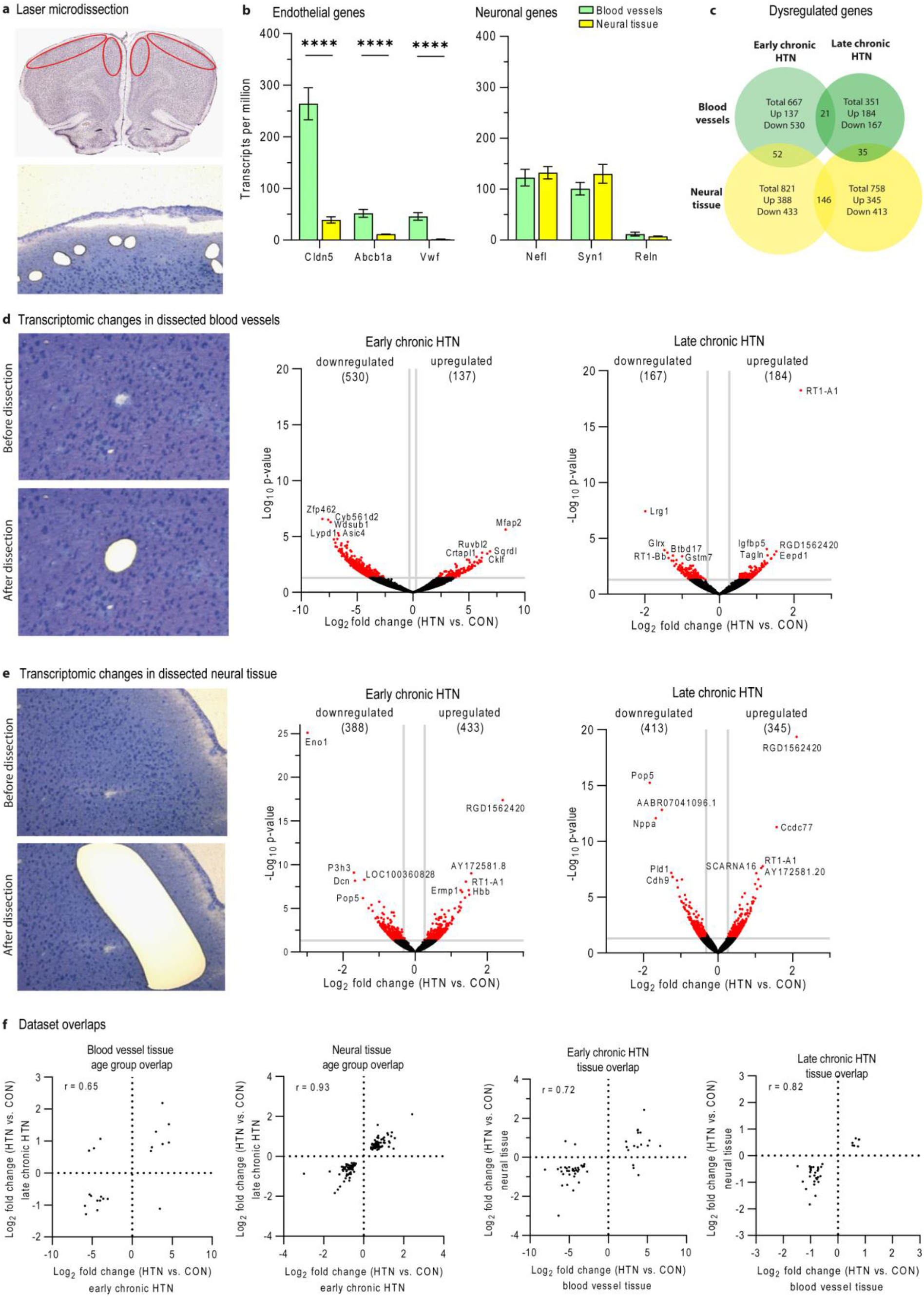
Validation of method and analysis of gene expression alterations. (**a**) Laser-microdissection of cortical blood vessels and neural tissue in toluene blue-stained, 15 µm thick, frontal brain slices. (**b**) Enriched expression of endothelial marker genes (Cldn5, Abcb1a, Vwf) in dissected blood vessels, but no enrichment of neuronal marker genes (Nefl, Syn1, Reln) in neural tissue, validating the enrichment of brain vascular cells in dissected blood vessels, containing also perivascular neurons. Significant group differences were assessed via paired, two-tailed Student’s t-test (details on further vascular cell types in **Supplementary Table 5**), and are highlighted via **** p < 0.0001; data are presented as mean ± s.e.m. (**c**) Number and overlaps of DEGs in early and late chronic hypertension in both tissues. (**d, e**) Exemplary field of views for blood vessel and neural tissue laser dissection. Volcano plots represent DEGs (p < 0.05 and fold change > 1.2) as red points and genes with no significant difference as black points. The x and y axes show log2 fold change and -log10 p-value, respectively. The top 5 down- and upregulated genes in each dataset are displayed. (**f**) Scatter plots depicting the logarithmic fold changes of genes. Pearson’s correlation was used to assess dataset overlaps. Abbreviations: Abcb1a, ATP binding cassette subfamily B member 1A; Cldn5, claudin 5; CON, control; DEG, differentially expressed gene; HTN, hypertension; Nefl, neurofilament light chain; Syt13, synaptotagmin 13; Tubb3, tubulin beta 3; Vwf, von Willebrand factor.

### Alterations in gene expression in chronic hypertension

To identify gene expression alterations in blood vessels and neural tissue in response to early and late chronic hypertension we selected differently expressed genes (DEG) based on a p-value < 0.05 and fold change greater than 1.2. We observed 667 and 351 DEGs in early and late chronic hypertensive blood vessels, respectively, and 821 and 758 DEGs in neural tissue of early and late chronic hypertension (***Figure 3c-e***). We identified sets of transcripts, that were dysregulated to a similar extent in the neural tissue and in blood vessels of both chronic hypertensive stages (***Figure 3f***). STRING interaction networks were created to visualize the physiological connections of all DEGs in each dataset (***Supplementary Figures 4-7***). Clusters of DEGs were associated with ribosomal function in blood vessels and neural tissue of early chronic hypertension. In addition, all four datasets showed clusters of genes involved in ubiquitination, indicating that intracellular proteostasis is altered. A human study on CSVD pathobiology in the frontal cortex also showed alterations in ubiquitination and protein catabolism [18].

### athobiology in early and late chronic hypertension based on GO function enrichment analysis

Functional sorting of significantly up- and downregulated GO-term and pathway enrichments revealed different regulatory patterns in early and late chronic hypertension (***Figure 4***). All DEGs corresponding to each of the GO-terms and pathways are displayed in ***Supplementary Table 6***.

**Figure 4.**
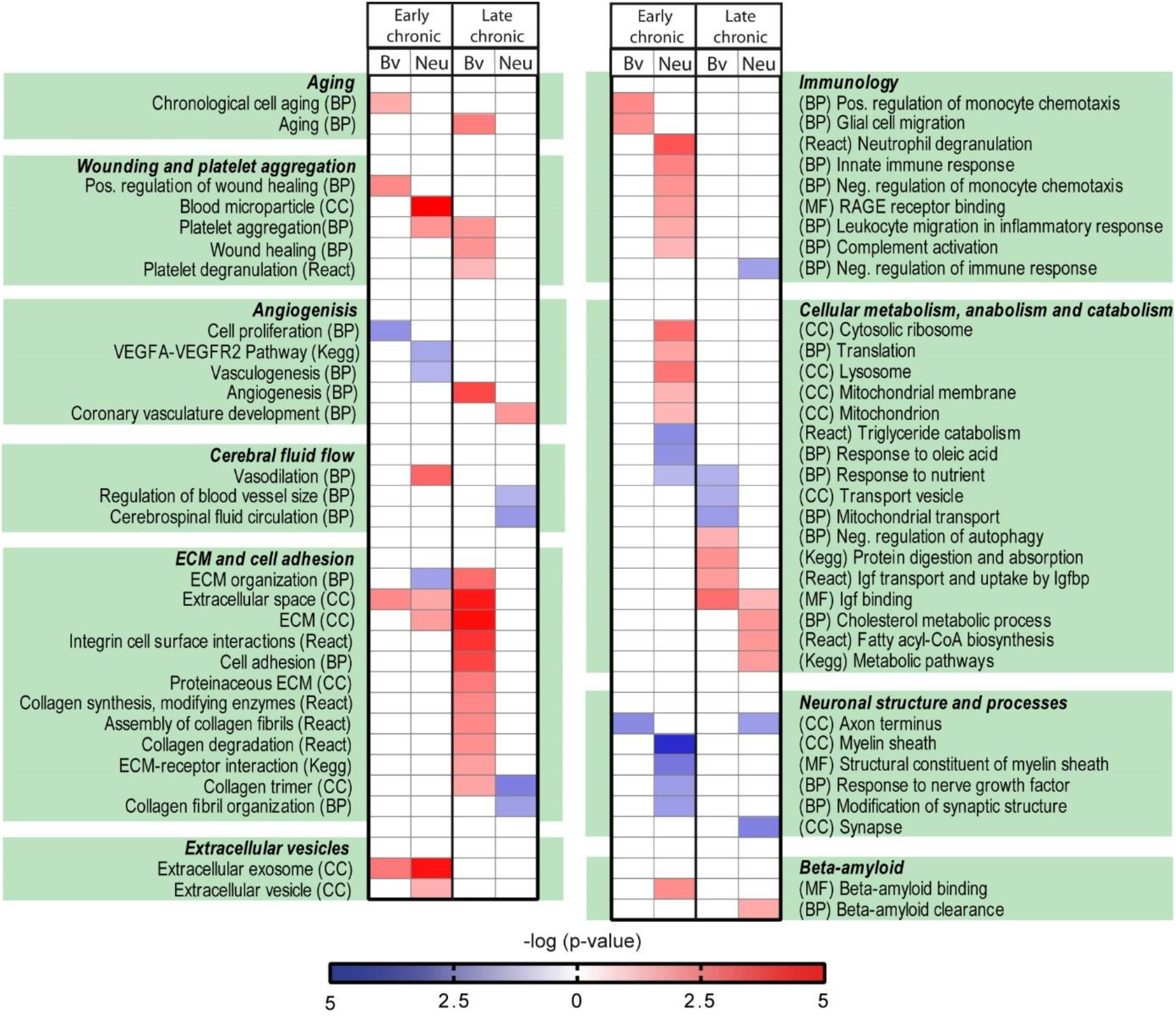
Pathobiology in early and late chronic hypertension based on GO-term enrichment analysis. Functionally sorted selection of GO-terms and pathways enriched among up- (red) and downregulated (blue) transcripts in early and late chronic hypertension were detected using the DAVID website. Color coding refers to -log (p-value). Categories of GO-term enrichments and pathways are displayed in brackets. Significance for GO-term enrichments was set using the EASE score with p-value < 0.05. The entirety of GO-terms, pathways and related genes can be seen in **Supplementary Table 6**. Abbreviations: BP, biological process; Bv, blood vessels; CC, cellular compartment; ECM, extracellular matrix; Kegg, Kegg pathway; MF, molecular function; Neu, neural tissue; React, Reactome pathway.

In line with strong plasma protein leakage in early chronic hypertension, increased transcript numbers associated with the GO-term blood microparticle were found in neural tissue, including extracellular globin transcripts *Hba-a2* and *Hbb*. Physiologically these transcripts are found in cell-free blood plasma [57] and are only able to enter the brain parenchyma when BBB integrity is impaired [58]. GO-term enrichments involving immunological processes were found in early chronic, but not in late chronic hypertension. An enrichment of upregulated transcripts including *C1qa, C1qc, S100a8/a9, Ccl2* and *Cklf* was associated with chemotaxis and immigration of innate immune cells. This molecular signature extends the leukocyte FACS results and indicates that peripheral immune cells exclusively participate early in chronic hypertension. Downregulated neuronal transcripts were associated with GO-terms of neuronal compartments, like axon terminus or myelin sheath and related neuronal functions, including response to nerve growth factor or modification of synaptic structure, indicating that neurons have an altered expression profile already in the early stages of chronic hypertension. Upregulated transcripts associated with GO-terms of cellular organelles and vesicles, including ribosomes, lysosomes, mitochondria and extracellular vesicles were found in neural tissue of early chronic hypertensive animals. These cellular processes, which are linked to protein synthesis, catabolism, energy demand and cell communication seem to undergo early alterations as well and might represent elevated cellular metabolic rates. A human microarray study in brain tissue from CSVD patients similarly showed upregulated genes associated with ribosomal function in the frontal cortex [18].

In late chronic hypertension blood vessels become stiff and the ECM accumulate, as demonstrated by a prominent pattern of GO-terms associated with ECM and adhesion molecules, possibly in an attempt to strengthen the vascular wall. Interestingly, enrichment of upregulated transcripts, including *Vegfb, Tgfa, Ctgf* and *Nov* was associated with angiogenesis in late chronic hypertension and further advances the molecular signature of increasing vessel density and endothelial cell proliferation. Additionally, transcripts associated with response to nutrients and genes associated with transport vesicles were downregulated in late chronic hypertensive blood vessels, potentially affecting adequate nutrient supply across the BBB. In cells contained in neural regions, genes related to metabolic pathways, fatty acid biosynthesis and cholesterol metabolism were upregulated, which may indicate an increased metabolic demand of the surrounding tissue. An enrichment of upregulated genes associated with inhibition of autophagy function, e.g. *Tbc1d14*, which is known to inhibit autophagosome formation [59], was also exclusively found in late chronic hypertensive blood vessels and might indicate dysregulated elimination of cellular waste. Transcripts with high Igf-1 binding affinity were upregulated in both tissues of this stage and might interfere with protective Igf-1 signaling.

Noticeable functional patterns were overlapping in both age groups and reflect continuous processes in chronic hypertension. An enrichment of upregulated transcripts associated with GO-terms of wound healing and accelerated aging were found in dissected blood vessels, suggesting that hypertension and related downstream signaling continuously create microinjuries and speed up vascular cell aging.

### Gene profile shift of vascular brain cells

Based on a molecular atlas [60], we studied which vascular brain cells displayed altered gene expression profiles in hypertensive blood vessels. The 50 most cell-specific genes for arterial smooth muscle cells (aSMC), FBs, endothelial cells and pericytes were screened for DEGs.

In aSMC-specific genes 11 out of 50 were found to be upregulated. They mainly participate in cell growth, differentiation, adhesion and the contractile system, pointing towards continuous aSMC remodeling, starting in early chronic hypertension. In contrast, 8 out of 50 FB-specific genes were upregulated almost exclusively in late chronic hypertension. All of them, except *Serping1*, function as ECM components or ECM binding proteins, stressing the importance to specifically recognize the role of FB in vascular stiffening and ECM remodeling. Transcriptional changes were also induced in endothelial cells, in that 6 out of 50 specific genes were dysregulated partially in early and partially in late chronic hypertension. These genes are linked to vascular activation, endothelial cell migration and proliferation. *Eng* is induced after vascular injury, during inflammation and is involved in angiogenesis [61]. *Podxl* is known to bind L-selectin on leukocytes, which could promote leukocyte immigration as well. Finally, 2 out of 50 pericyte-specific genes were found to be dysregulated in chronic hypertensive blood vessels, namely downregulated *Ecm2* in early chronic and upregulated *Rgs5* in late chronic hypertension, of which the latter is a known vascular detachment marker of pericytes [62, 63] (***Figure 5***).

**Figure 5.**
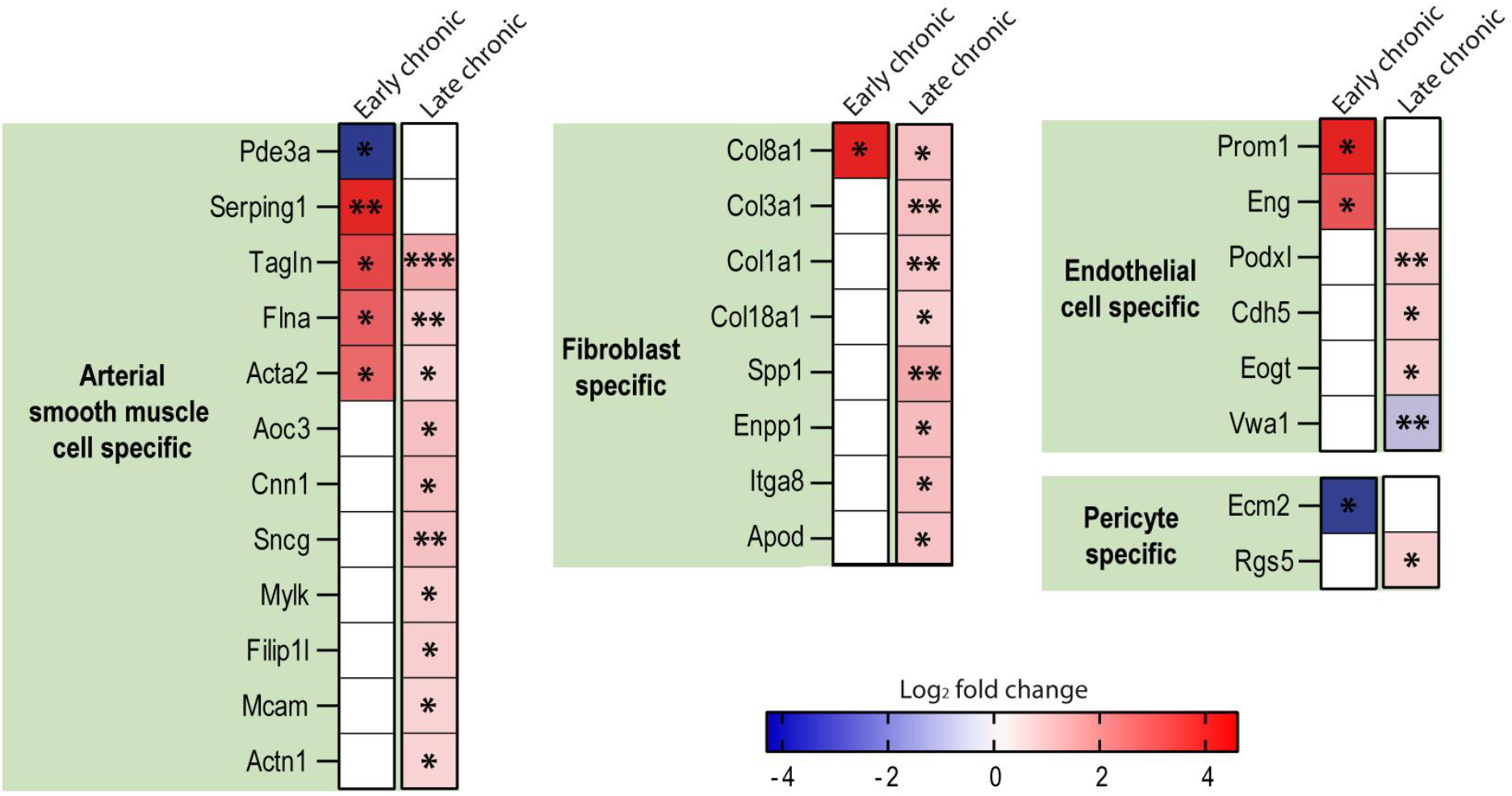
Vascular cell-specific DEGs in blood vessels of early and late chronic hypertension. Logarithmic fold changes are presented, in which red indicates upregulation and blue downregulation of the respective transcript in chronic hypertension compared to age-matched controls. Level of significance is displayed as *p < 0.05, ** p < 0.01 and ***p < 0.001. Abbreviations: DEG, differentially expressed gene.

### ^18^F-FDG PET reveals cortical hypermetabolism in early chronic hypertension

The upregulated transcripts of cellular metabolism and inflammatory processes prompted us to screen for alterations in brain glucose-metabolism using ^18^F-FDG PET. We used a standard protocol of ^18^F-FDG PET imaging based on a 45 min uptake phase, followed by read-outs of the distribution of the trapped, phosphorylated tracer in one static scan. We noted increases of up to 30% in early chronic hypertensive animals compared to age-matched controls. The cingulate and retrosplenial cortices as well as motor and somatosensory cortices were mostly affected (***Figure 6***). In late chronic hypertension, the spatial distribution of affected regions is largely similar but the effect size is markedly reduced as compared to early chronic hypertension.

**Figure 6.**
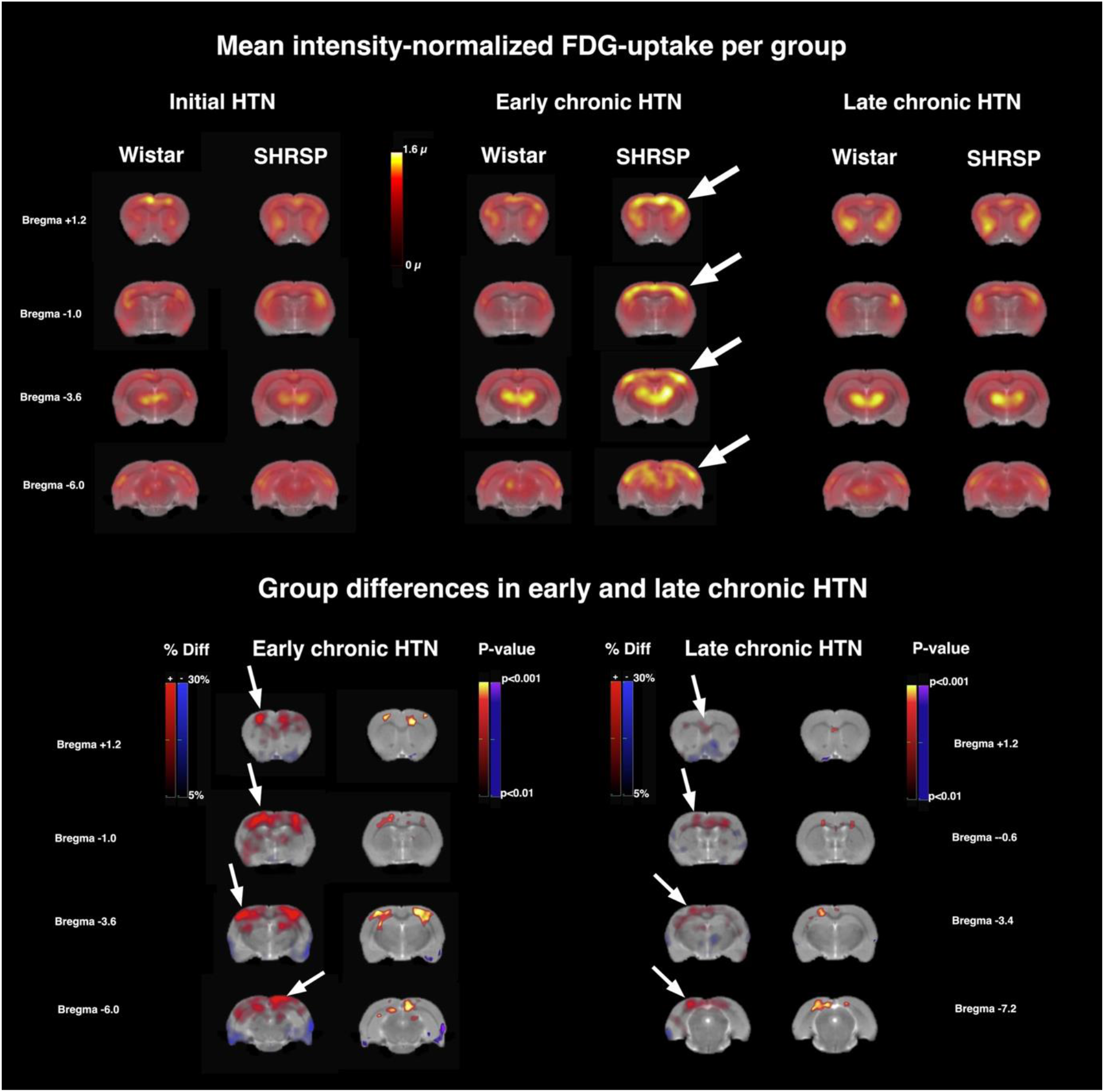
Brain regional glucose metabolism studied with ^18^F-FDG PET in chronic hypertension. Sections of group-mean intensity-normalized 18F-FDG uptake are shown in the upper half of the figure, overlaid on an anatomical reference MRI. Color scale is in multiples of the global mean µ. In the lower half, percentage differences and statistically significant differences between early and late chronic hypertensive rats and age-matched Wistar control rats are shown. Warm colors indicate higher, blue colors lower metabolism in hypertensive rats. Note the marked increase in metabolism in early chronic HTN of up to ca. 30% in cortical areas (arrows) including motor and cingulate cortex (Bregma +1.2; -1.0) and in parts of the hippocampus. In late chronic HTN the spatial distribution of affected regions is largely similar but the effect size is markedly reduced as compared to early chronic HTN. Animal numbers: n = 4 CON vs. n = 6 initial HTN, n = 6 CON vs. n = 8 early chronic HTN, n = 5 CON vs. n = 6 late chronic HTN. Abbreviations: CON, control; ^18^F-FDG-PET, ^18^F-Fluorodeoxyglucose positron emission tomography; HTN, hypertension; MRI, magnetic resonance imaging; SHRSP, spontaneously hypertensive stroke-prone rat.

## Discussion

We applied a variety of methods to understand pathomechanisms and their transcriptomic signatures in different stages of arterial hypertension in the rat frontal brain. We observed (i) frontal brain-specific behavioral changes and cognitive impairments as a consequence of chronic hypertension, (ii) characteristic microvascular pathologies, (iii) fast recruitment of activated microglia to the blood vessels, early immigration of peripheral immune cells, (iv) an energy-demanding hypermetabolic state and (v) vascular adaptation mechanisms in late chronic stages, including angiogenesis and vessel wall strengthening by upregulation of cellular adhesion molecules and extracellular matrix. (vi) Additionally, we identified late chronic vascular accumulation of Igfbp-5 in the brains of hypertensive rats, a signature of Alzheimer’s disease, attenuating protective Igf-1 signaling.

### Behavioral changes and cognitive impairments

Reduced anxiety, as measured by time in the center of an open field and deficits in inhibitory sensorimotor gating, as determined by PPI of an ASR, are consistent with previous studies in chronic hypertensive rat models [64–66], but have not yet been linked to behavior in initial hypertensive stages. Here, we show that these alterations occur early and persist in chronic hypertension. Cortical neuronal networks of the frontal lobe are significantly involved in PPI. In humans alterations in PPI are linked to altered focused attention and processing speed, mimicking typical impaired cognitive domains in CSVD [27, 55, 67]. Thereby PPI might be indicative of altered neuronal network functions in chronic hypertension. Reduced activity in the open field test was observed in all age groups and thus did not reflect specifically chronic hypertension-related behavioral changes, but is in line with results from an angiotensin II-induced hypertensive mouse model [68]. In contrast normotensive ATP11b knockout rats similarly show reduced activity and it is known that the ATP11b gene has a homozygous exonic deletion mutation in SHRSP, but not in Wistar control rats, which also indicates a genetic contribution [69].

### Microvascular pathology

Hypertension-related microvascular pathology comprised CMB and EPVS formation, worsening with prolonged hypertension. Cumulative data from a large number of rodents were retrospectively used to compute a robust regression model on whole brain CMB prevalence probability, which was overall tripled in hypertensive rats. Human MRI studies, that prospectively compared CMB development between normotensive and hypertensive individuals, are consistent with this finding and showed a threefold increased odds ratio in hypertensive patients [70, 71]. The frequency of EPVS in the frontal cortex was already increased in initial hypertension and worsened in chronic stages, also reflecting findings of a recent meta-analysis of human MRI studies, in which hypertension and age favored EPVS formation [72]. BBB leakage first occurred in early chronic hypertension with transcriptomic signatures of blood microparticles in the brain tissue and was partially recovered in late chronic hypertension, accompanied by an upregulation of vascular ECM and cell adhesion molecules (Cdh2, Cdh5). These proteins are known to promote BBB integrity [73–75] and therefore may reflect molecular compensatory mechanisms to reduce BBB leakage. Studies that assessed BBB leakage individually in early and late chronic hypertensive rats comparably to the here investigated age groups showed similar biphasic results [76–78]. In 17 months old hypertensive rat brains, cortical BBB leakage was increased again compared to normotensive controls [79]. One could hypothesize that BBB leakage appears early in the disease continuum, is then partially compensated and reappears when compensation mechanisms fail. A recent study described a core BBB dysfunction module, comprising n = 54 upregulated genes in four different neurological diseases [80]. Comparing this list with our vascular datasets we identified n = 4 overlaps in early chronic hypertension (*Adamts8, Ccl2, Serping1, Tubb6*) and n = 4 overlaps (*Col3a1, Plekho1, Spp1, Igfbp5*) in late chronic hypertension. This overlap verifies signatures of BBB dysfunction in both chronic stages and we speculate that the mechanisms underlying this regulatory pattern are of general relevance and might be a therapeutic target to mitigate BBB dysfunction, independent of the underlying disease. Human data on BBB leakage are difficult to compare due to heterogeneous cohorts and different methodologies. Neuropathological and MRI studies showed an increase in BBB leakage due to chronic hypertension, but this rather reflects end-stage CSVD with inclusion of stroke and dementia cases [81– 84]. Conversely, cross-sectional cerebrospinal fluid (CSF) studies in older adults found no influence of hypertension on the CSF / serum albumin ratio [85, 86]. To better characterize the important role and dynamics of BBB leakage in chronic hypertension prospective studies should be preferred. They should be designed with the potential to capture the stage-dependent and possibly reversible nature of BBB leakage on an individual basis. Additionally, the disease duration and treatment effectiveness of BP-lowering medications as well as the personalized intake of different types of antihypertensives should be integrated into these analyses instead of only dichotomizing into hypertensive / not hypertensive.

### Angiogenesis

Brain function depends on an adequate supply of nutrients and oxygen, provided by a dense microvascular network. The microvasculature in the aging and hypertensive frontal rat cortex exhibited high plasticity. In late chronic hypertension, increased microvascular density and elevated expression of proliferative and angiogenic markers were signs of active angiogenesis. This is in line with previous rat studies that found angiogenic responses in chronic hypertension [69, 78, 87, 88]. We previously showed that cortical blood flow is reduced in chronic hypertensive rats [11], similar to a recent large human MRI study that assessed cerebral blood flow [89]. Angiogenesis might thereby be a compensatory mechanism for insufficient tissue supply. This concept is well-described for other human neurological pathologies with reduced blood flow, such as in and around cortical microinfarcts or in neurodegenerative diseases such as AD [90–93]. However, these newly formed vessels are suspected to be dysfunctional with increased BBB permeability, thrombogenic potential and disturbances in microcirculation [90–93]. A recent human neuropathological study supports this hypothesis. Early-stage hypertensive CSVD patients showed a trend towards an increased vessel density in the frontal white matter compared to controls. However, the microvasculature showed frequent string vessels, which correspond to collapsed basement scaffolds connected to the capillary network, often lack endothelial staining and indicate endothelial cell damage or even apoptosis [82]. Microvascular rarefication then likely occurs in a late disease stage as a consequence of this endothelial cell apoptosis [6].

### Early microglial activation and immune cell infiltration

It is widely accepted, that neuroinflammation plays a pivotal role in chronic hypertension and microvascular pathologies. It is however insufficiently understood in which disease stage resident and peripheral immune cells become involved and which molecular signaling pathways have to be considered. We here show microglial recruitment and activation already in initial hypertension and in absence of BBB leakage, followed by reduced responsiveness to BBB leakage in chronic stages of hypertension. In human AD there is a concept of biphasic microglial activation within the course of disease, in which the first beneficial activation phase is induced by accumulating Aβ, followed by a phase of exhaustion and decreased activation, which finally results in an overshooting neurotoxic inflammatory response [94]. One might speculate that initial hypertension represents a similar activating trigger, followed by a period of habituation to reduce neuroinflammation. The resulting chronic low-grade microglial activation might lead to cellular exhaustion, which limits responsiveness to vascular injuries in late stages of the disease. In contrast, cortical infiltration of peripheral immune cells and their inflammatory transcriptomic signatures were observed when BBB leakage was most severe, potentially facilitating the crossing of the BBB. Chemotactic signaling via Ccl2 from vascular cells recruited leukocytes to the site of injury, which is in line with human data, where increased levels of Ccl2 were found in blood samples of hypertensive patients, and correlated with the degree of hypertension-associated organ damage [95, 96]. The upregulation of C1q transcripts in the neural tissue of hypertensive rats points to an involvement of the complement signaling system and might be a link to several downregulated neuronal and synaptic transcripts, since it has been shown that the elimination of synapses is driven in a C1q-dependent manner [97, 98].

### Increased glucose consumption in chronic hypertension

To the best of our knowledge ^18^F-FDG-PET brain imaging has not been performed in hypertensive rats so far. We used a standard protocol with an acquisition of one static image 45 min after injection. During the long tracer uptake time non-phosphorylated ^18^F-FDG moves out of the brain and therefore differences in tracer-content represent differences in glucose-phosphorylation rather than glucose-transport rates or blood flow [99]. We here show highly increased glucose consumption in early chronic hypertension, which was still present in the late chronic stage, but to a markedly reduced extent. In the humans brain, glucose consumption decreases during aging [100] but complex regional alterations in glucose consumption have been described in the spectrum of AD. Neuronal activity and glucose consumption show an inverted u-shape in the disease course of AD. While end-stage AD is typically accompanied by neuronal/synaptic hypoactivity and temporoparietal hypometabolism, particularly in early and silent stages, neuronal circuits are hyperactive and regionally increased glucose-consumption has been found [101–104]. In principle, these increased glucose consumption rates could be due to increased neuronal activity. Recent studies in both patients and animal models of dementia point also to activated microglia driving glucose hypermetabolism [105, 106], which is in line increased metabolic demands after experimental stroke and intracerebral hemorrhage (ICH), in which activated microglia and infiltrating inflammatory peripheral immune cells contribute to up to 50% of increased glucose consumption [107, 108]. The findings are discussed controversially, since activated microglia represent a rather small proportion of cells compared to the total number of cells in a volume-element in FDG-PET imaging and additionally reactive astrocytes have been suggested to play a role in elevated glucose consumption [109, 110]. Oligodendrocytes could also contribute to an increase in energetic demands. A multi-tracer PET imaging study showed a shift from mitochondrial oxidative phosphorylation to aerobic glycolysis in non-lesional white matter of CSVD patients compared to age-matched controls [111]. These findings are concordant with experimental studies in mice with mitochondrial respiration deficiency, showing that elevated glial aerobic glycolysis is compensatory to provide trophic support to neurons [112]. Similar mechanisms are conceivable in the context of hypertension if cellular energy production shifts from mitochondrial respiration to aerobic glycolysis, which is much more glucose-consuming. We, therefore, propose that neuronal hyperactivity, inflammatory signatures and alterations in energy production are related to increased metabolic demand in the silent phase of chronic hypertension. We here provide evidence for the first time, that brain energy metabolism is remarkably altered in chronic hypertension. Future studies have to elucidate, which cell types, cellular mechanisms and energetic metabolites are involved in this metabolic shift and how it affects cellular function. FDG-PET in combination with measures of inflammation and neuronal activity might be a new and non-invasive opportunity to investigate altered brain metabolism in the early and silent stage of chronic hypertension in humans.

### Igfbp-5 as a potential new candidate in disease pathophysiology

Igf-1 is a trophic factor for vascular as well as neuronal cells whose growth-promoting actions are mediated via the Igf-1R. Expression rates of Igf-1R in the brain is highest in endothelial cells and neurons [113]. Circulating Igf-1 crosses the BBB by luminal endothelial Igf-1R binding and is transported to the basolateral side of the vessel after which it is released and available for adjacent neurons. This process is even enhanced in areas of high neuronal activity and thereby provides neurotrophic factors on demand [114]. High Igf-1/Igfbp-5 and Igf-1/Igfbp-2 binding affinity is known to reduce Igf-1R activation and downstream signaling, when these binding proteins accumulate [115–117]. We identified highly upregulated *Igfbp-2* and *Igfbp-5* in hypertensive blood vessels and neural tissue from rats and prominent accumulation of Igfbp-5 in the frontal white matter of hypertensive CSVD patients, which potentially reduces Igf-1 signaling for vascular cells and adjacent neurons. In humans, reduced circulating Igf-1 is associated with endothelial and SMC dysfunction, vasoconstriction and increased risk of hypertension-induced microvascular pathologies or stroke [6, 118, 119] and thereby reflects a marker of poor vascular health. Large proteomic aging studies identified high Igfbp-5 protein levels in frontal cortical areas to be associated with faster decline in cognitive and motor function prior death. This association was partially explained by neuropathological markers of AD and frontal white matter microstructural changes but it was suggested that Igfbp-5 is also mediating the effects of other yet unmeasured neuropathologies [120–123]. Elevated expression of Igfbp-5 resulted in motor neuron degeneration and myelination defects due to insufficient trophic support of axons in human diabetic neuropathy as well as in Igfbp-5 overexpressing rodents [115]. One could hypothesize that similar effects occur locally in the white matter of chronic hypertensive individuals, which would link the susceptibility of frontal white matter tracts to chronic hypertension [17, 19–21]. Igf-1 signaling also promotes Aβ clearance through the BBB and is thus involved in protecting against AD pathology [124]. However, Igfbp-5 is an upregulated BBB dysfunction signature gene, even for months after the initial event. This could inhibit the protective Igf-1 effects against AD pathology and might help to explain the increased risk for chronic hypertensive individuals to develop AD.

### Chronic hypertension as a risk factor for stroke and AD

Hypertension and age are the most relevant risk factors for vascular pathologies in the brain, which lead to disability and cognitive decline [125, 126]. A recent meta-analysis summarized genetic variants and risk genes that are associated with the development of lacunar stroke and ICH. Out of 16 risk genes for lacunar stroke we identified n = 3 DEGs in hypertensive blood vessels (*Camk2d, Zdhhc20, Fam117b*) and n = 2 DEGs in hypertensive neural tissue (*Camk2d, Epb41l3*). Out of 12 risk genes for ICH n = 1 was differentially expressed in hypertensive blood vessels (*Wdr12*) and n = 1 in hypertensive neural tissue (*Apoe*). Gene candidates that are involved in the development of vascular pathologies based on genetic intrinsic and environmental extrinsic factors could be of general relevance in the disease pathophysiology and thereby interesting new candidates for future research.

The contribution of vascular risk factors to neurodegenerative diseases such as AD is gaining research interest as well. Genome-wide association studies have nominated genes that contribute to AD risk and pathophysiology. We wondered whether chronic hypertension has an impact on expression of these candidate genes and thereby potentially contributes to increased risk of disease development. Out of the top 45 AD risk genes [127] we identified n = 1 DEG in hypertensive blood vessels (*Bin1*) and n = 6 DEGs in the hypertensive neural tissue (*Apoe, Bin1, Inpp5d, Abi3, Picalm, Adamts4*). Single-cell RNAseq revealed that all of these overlapping genes, except Adamts4, are strongly expressed by microglia compared to other cell types [113, 127], indicating that microglial gene expression perturbations in chronic hypertension could contribute to increased risk of AD development. We further investigated whether blood vessels in chronic hypertension and human AD share similar transcriptomic signatures. Comparing our datasets with n = 463 DEGs from blood vessels in AD [127] we identified n = 8 overlaps in early chronic hypertension (*Trio, Cmss1, Map2k6, Btbd9, Pde3a, Pdzd2, Pias2, Hivep1*) and n = 6 overlaps in late chronic hypertension (*Itga8, Plxnc1, Ptges3, Emcn, Cdh2, Mfsd2a*). Interestingly all overlapping DEGs in the early phase were downregulated and all in the late phase were upregulated in both conditions. In summary, these candidate genes might serve as a molecular target for microglia and vascular cell perturbations, which could partially explain the contribution of hypertension to AD risk.

### Limitations

There are several limitations to this study. Most of our experiments focused on frontal cortical areas, which are known predilections for hypertension-related alterations. However, the generalizability of stage-dependent mechanism to other brain regions is partially limited. To overcome this issue, we performed microvessel isolation and FACS analysis in the entire cortex and ^18^FDG-PET imaging in the entire brain, which made our major findings transferable to the whole cortex. Future studies might also include the hippocampus as a second interesting target region. Although different experimental models show similar pathomechanisms in hypertension [68, 78], their stage-dependent nature should be validated in these different models as well (e.g. spontaneously hypertensive rats or angiotensin II-induced hypertensive mice) and particularly in the human brain of hypertensive individuals to generalize the concept and rule out genetic contributions in the SHRSP model [16, 69]. Lastly the integration of our results into human literature remains difficult since longitudinal studies on an individual bases are sparse [5, 13] and neuropathological studies often address advanced and end-stages of the disease [128]. This conflict can be overcome if future research focuses on characterizing the silent phase of hypertension, before structural pathologies occur. Clinical studies should integrate disease duration and follow up on individuals after the diagnosis of hypertension and autopsy studies might focus on younger individuals or well-prepared study designs with longitudinal characterized subjects comparable to the Lothian birth cohort.

## Acknowledgements

This work was funded by German Research Foundation (GRK SynAge 2413, TP7 to S.S. and A.D.; SRC 1436 Neural Resources of Cognition, A05 to A.D.; 9^th^ Nachwuchsakademie Medizintechnik MA 9235/1-1 to H.M.), by Medical Faculty of the Otto-von-Guericke University Magdeburg (scholarship to P.U.) and by Federal state Saxony-Anhalt and the European Structural and Investment Funds.

## Author contributions

PU, LM, MB, NL and SJ performed most of the experiments. SJ did behavioral tests with help from AB; PU performed immunofluorescence experiments and image analysis; MH, SJ and PU did histological examinations; LM performed microvessel isolation, PCR and FACS and NL did laser dissection, RNAseq and PCR with help from MB. HM, MH, MK, MW and SA helped with image and data analysis. DG, MT, PB, TK and AO were involved in FDG-PET experiments and JG did the FDG-PET data analysis. DY-H performed the human experiments. PU did most of the data visualization. MS, ID and SS advised and directed the experiments and were involved in the conceptualization of the study. PU mostly wrote the paper, with input from all authors. SM, H-JH, GD-V, SV and AD. Reviewed and edited the manuscript and contributed to the interpretation of the data. All authors approved the version to be published

